# Predicting the principal components of cortical morphological variables

**DOI:** 10.1101/2022.07.07.499214

**Authors:** V. B. B. Mello, F. H. de Moraes, B. Mota

## Abstract

The generating mechanism for the gyrification of the mammalian cerebral cortex remains a central open question in neuroscience. Although many models have been proposed over the years, very few were able to provide empirically testable predictions. In this paper, we assume a model in which the cortex folds for all species of mammals according to a simple mechanism of effective free energy minimization of a growing self-avoiding surface subjected to inhomogeneous bulk stresses, to derive a new set of summary morphological variables that capture the most salient aspects of cortical shape and size. In terms of these new variables, we seek to understand the variance present in two morphometric datasets: a human MRI harmonized multi-site dataset comprised by 3324 healthy controls (CTL) from 4 to 96 years old and a collection of different mammalian cortices with morphological measurements extracted manually. This is done using a standard Principal Component Analysis (PCA) of the cortical morphometric space. We prove there is a remarkable coincidence (typically less than 8^*◦*^) between the resulting principal components vectors in each datasets and the directions corresponding to the new variables. This shows that the new, theoretically-derived variables are a set of natural and independent morphometrics with which to express cortical shape and size.

## 1 Introduction

Why the cerebral cortex folds in some mammals is a fundamental question in neuroscience that still lacks a broadly accepted mechanism. It is reasonable to think about the cortical folding as a characteristic fixed by natural selection because it allows a dramatic increases in cortical size and complexity, while preserving the cranial size, axonal elongation and metabolic cost [31, 32]. However, the evolutionary argument is not sufficient to unravel the full process that results in the surface folding. Empirical evidence points towards an integrative explanation [18], unifying bio-mechanical, genetic and cellular signaling mechanism.

From a purely bio-mechanical point of view, there is a variety of models proposed to explain the physics of cortical folding: buckling from a constrained expansion [33, 34], warping due to differential tangential expansion [28] or emerging from axonal elongation [13]. Despite the qualitative compatibility with the cortical morphology, all these theories failed to provide a set of empirical testable predictions to further investigate their validity.

Following the bio-mechanical approach, [22] proposed a folding model hypothesizing that the cortex is a crumpled elastic surface in thermal equilibrium (like a crumpled paper ball) [15]. The model is based on the physics of minimization of effective free energy of a growing self-avoiding surface subject to inhomogeneous bulk stresses. It predicts a power-law relationship between cortical thickness (T), exposed (A_*E*_), and total area (A_*T*_) (Equation 1)

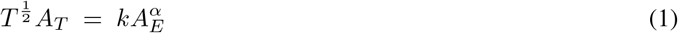

for which the exponent *α i*s a universal constant with a theoretical value of 1.25 and *k i*s a dimensionless parameter related to the mechanical properties of cortical tissue. The value of *k n*ot specified by the model, but is apparently highly conserved across various mammalian orders. The most immediate consequence of this model is that the usual set of geometrical variables {*T*^2^, *A*_*T*_, *A*_*E*_} used to describe the cortex should not be independent, but rather correlated in a highly restrictive way. Representing these variables in the {log *T*^2^, log *A*_*T*_, log *A*_*E*_} space so that products of power laws become linear combinations of the log-variables, one can define

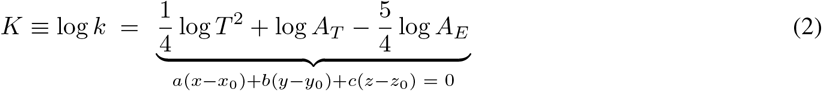

In this log-space, Equation (1) can be re-interpreted as an assertion that actual cortices must lie at or near a *K = constant* plane, called henceforth the theoretical cortical plane. Thus, as long as it is constrained to a plane perpendicular to the vector 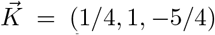 in log-space, cortical morphology should otherwise be compatible with any set of values {*T*^2^, *A*_*T*_, *A*_*E*_}. Taking a step further, Equation 1 is a power-law relation between the intrinsic and extrinsic values for cortical areas, measured in unites of squared cortical thickness, suggesting that the cortex is self-similar over some range of length scales, i.e, approximately fractal with fractal dimension *f*_dim_ = 2*α*.

The predictions of this simple statistical physics model were experimentally tested in a variety of studies. Firstly, it was found that for 63 different mammalian species cortical folding scales universally according Equation 1 with *α*_exp_ = 1.31 *±* 0.01 (when considering a heterogeneous, i.e, multiple acquisition protocols dataset) [22]. This work was extended in [40], where it was shown that not only the universal law also holds within a single species MRI dataset with *α*_exp_ = 1.25 *±* 0.01 compatible with theoretical predictions, but also provided evidence that a deviation from the universal rule could be used to identify diseases. The discrepancy between the observed *α* and the theoretical prediction for the comparative neuroanatomy study needs further investigation to determine if it is due to a systematic effect on the heterogeneous dataset or more profound information about the model’s limitations. Then, in [23], it was observed that the white matter volume and white/gray matter ratio in mammalian species are a consequence of the universal scaling of cortical folding [42]. The first evidence of the self-similarity of the cortical surface was presented in [41] with experimental verification of the universal scaling law across lobes of the same cortex. More recently, [37] provided the first proof by construction of a universal archetypal fractal primate cortical shape, of which all actual cortices are approximations of a specific range of scales.

As a consequence, and regardless of the validity of the initial model, it is clear that the set of geometrical variables {*T*^2^, *A*_*T*_, *A*_*E*_} is indeed highly correlated. In [38], it was explicitly shown that by representing these variables in the {log *T*^2^, log *A*_*T*_, log *A*_*E*_} space one can define K (Equation 2).

In log space, healthy human cortices lie at or near the theoretical cortical plane, defined by points for which *K = constant*, in agreement with the crumpled cortex model. In an attempt to understand the differences between cortices at or near the cortical plane, [38] derived a set of orthogonal morphological variables, starting with the nearly invariant *K* 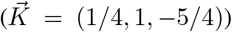 that is used to define the plane itself. Since *k i*s dimensionless, it is not affected by an isometric scaling log *T*^2^ *→* log *λT*^2^, log *A*_*T*_ *→* log *λA*_*T*_ and log *A*_*E*_ *→* log *λA*_*E*_, in which *λ i*s a positive real number; thus one can define both a direction 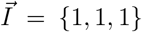 that correspond to this isometric scaling, and a corresponding new variable *I =* log *T*^2^ + log *A*_*T*_ + log *A*_*E*_ that captures all information about cortical size. The third and last vector to both 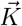 and 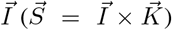, and is associated with the dimensionless variable 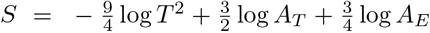 that would captures most of the variation in shape across actual cortices. These new variables are plausibly a natural way to decompose morphology in terms of its size; and its variant, and invariant shape components. Empirically, these new variables were shown to better discriminate between typical and atypical structural changes, like aging and diseases [38].

The existence of the cortical plane is by itself a significant empirical result for neuroscience. In principle, cortical morphology could be biologically compatible with any set of values {log *T*^2^, log *A*_*T*_, log *A*_*E*_}. In practice, it has been shown to be restricted to a plane. From a more instrumental perspective, this result implies that: (i) using the usual morphological variables without taking into account their correlations would severely distort statistical inference; (ii) there is clearly a direction in the log-space with the least morphological variation between cortices that is very close to the direction defined by the vector 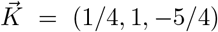, as predicted by the crumpled cortex theory; (iii) diseases that affect cortical morphology could be characterized by particular deviations from and along the cortical plane in relation to healthy controls. It is also important to emphasize that although the variables *K, S* and *I w*ere derived from a specific model, their use to express and analyze morphology over the cortical plane is model-agnostic.

The objective of this work is to test and systematize the description of cortical morphology, contrasting the old and new variables in terms of their statistical interdependence. To test the independence between the new variable set {*K, S, I}* we performed a standard Principal Component Analysis (PCA) to obtain the independent components of variance in the {log *T*^2^, log *A*_*T*_, log *A*_*E*_} space, for two different cases: across multiple species and across healthy human subjects (obtained from [3, 22]). The principal components (PC) expected for the first case are the independent set {*K, S, I}* [41]. For humans, in addition to a nearly invariant *K*, it is known that the average cortical thickness varies very little in comparison to *A*_*T*_ and *A*_*E*_, a direct consequence of being the same species. With the extra nearly invariant direction log *T*^2^, one would expect that the component with the most variance should lie along the 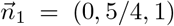 direction. The remaining PC can then be directly found by the cross product 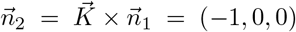. The PC vectors obtained from the data were then directly compared to the theoretical vectors 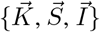 and a new set of vectors 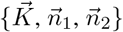 derived in this work to describe a single species.

To test the statistical robustness of our results, a simulated dataset was generated according to Equation 1 to address the sensitivity of the PCA technique to the dataset’s true principal directions. Finally, our results were validated by an alternative method. Rewriting Equation 1 as

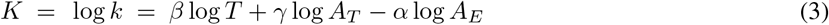

one realizes that the model prediction are actually the exponents *α =* 5/4, *β =* 1/2 and *γ =* 1, and that obtaining the least variance PC corresponds to performing a joint fit of these parameters. So, the same task was performed with a standard maximum likelihood approach combining information from both humans and multiple species datasets.

## 2 Results

### 2.1 Human MRI multi-site harmonized dataset

Performing the PCA in the human harmonized dataset [1, 3, 9, 19, 25, 30, 40, 41], one obtain the results summarized in Figure 1. We analyzed separately subjects in the 20-40 yo age range, who presumably have a fully developed cortex (in broad terms, after development but before significant aging), and subjects in the age range between 20-60 years, for whom age effects were mathematically removed (henceforth “de-aged”) [3]. Table 1 summarizes the differences between the principal components and the theoretical expectation. These differences do not change substantially between the two groups (see Appendix 5).

**Figure 1:**
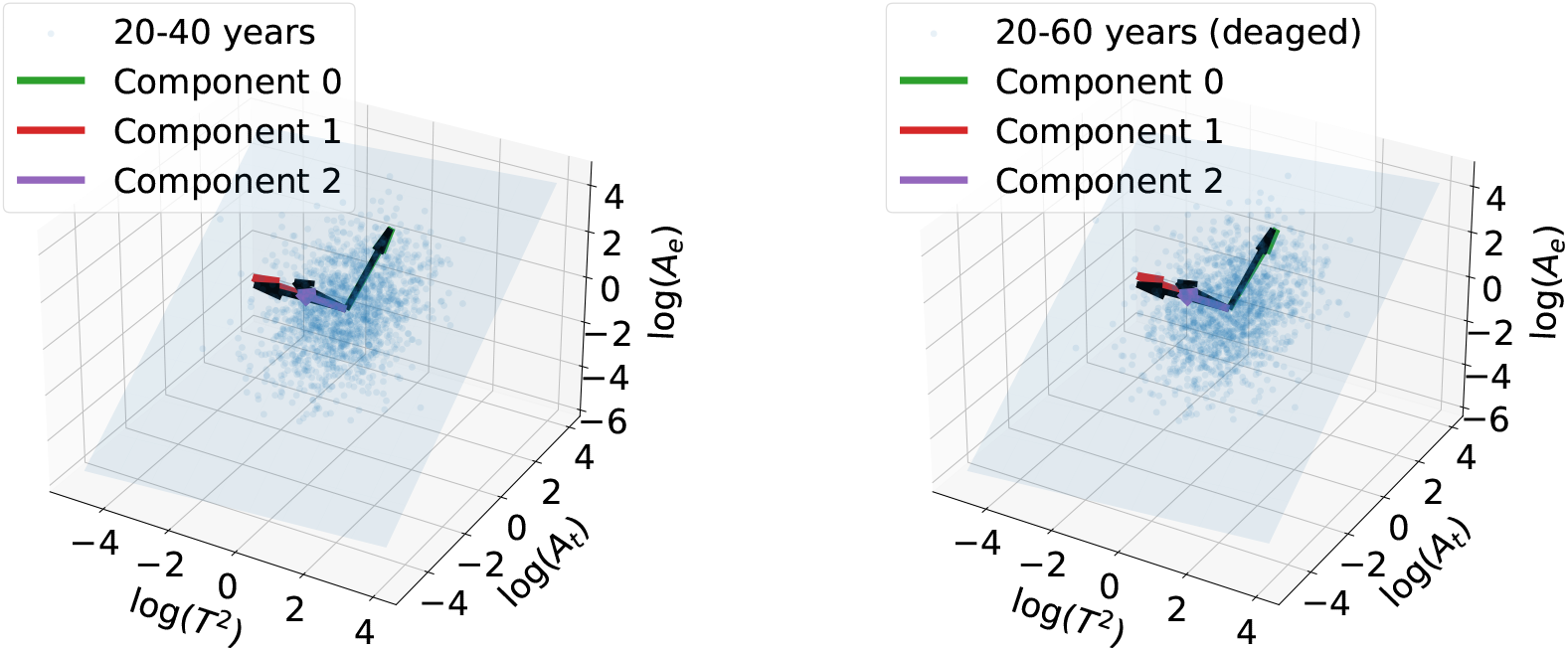
Cortical plane for the human MRI dataset and the vectors reconstructed by PCA. On the left panel, we considered only subjects in the age range of 20-40 years. On the right panel, the de-ageing procedure was applied, allowing the analysis of the full age range in which the model was experimentally validated (20 to 60 years old). The results of the two approaches are consistent. The arrows in black represent the principal components predicted by the model. The angular difference between theoretical components and the experimental ones are *θ*_diff_ *< 8*^*◦*^. The results are in remarkable agreement with the theoretical predictions.

**Table 1:** The normalized vectors found by PCA and their respective deviation from theoretical expectation for the MRI humans dataset (single species). The components are ordered by the Explained Variance Ratio (EVR).

| Dataset | Direction | Theoretical | PCA | $\theta_{\text{diff}}$ | EVR |
| --- | --- | --- | --- | --- | --- |
| 20-40 years | $\hat{v}$ | $\hat{n}_1 = (0, 0.781, 0.625)$ | $(0.056, 0.704, 0.707)$ | $7.9^\circ$ | 0.65 |
| | $\hat{u}$ | $\hat{n}_2 = (-1, 0, 0)$ | $(-0.996, 0.089, -0.009)$ | $6.9^\circ$ | 0.33 |
| | $\hat{w}$ | $\hat{K} = (-0.154, -0.617, 0.771)$ | $(-0.070, -0.704, 0.707)$ | $7.5^\circ$ | 0.02 |
| 20-60 years<br>(deaged) | $\hat{v}$ | $\hat{n}_1 = (0, 0.781, 0.625)$ | $(0.077, 0.703, 0.707)$ | $7.9^\circ$ | 0.65 |
| | $\hat{u}$ | $\hat{n}_2 = (-1, 0, 0)$ | $(-0.994, 0.110, -0.0004)$ | $6.9^\circ$ | 0.33 |
| | $\hat{w}$ | $\hat{K} = (-0.154, -0.617, 0.771)$ | $(-0.078, -0.703, 0.707)$ | $7.5^\circ$ | 0.02 |

### 2.2 Comparative Neuroanatomy

The original work by Mota & Herculano-Houzel included two primary datasets of mammals, analyzed separately here: Herculano-Houzel’s own data (referred here as dataset A) [10, 11, 16, 24, 27, 35], and previously published cortical morphological measurements (dataset B) [2, 5, 12, 20, 26]. Regarding the comparative neuroanatomy dataset, the PCA result is shown in Figure 2. The deviation between the principal components and the theoretical predictions are summarized in Table 2.

**Figure 2:**
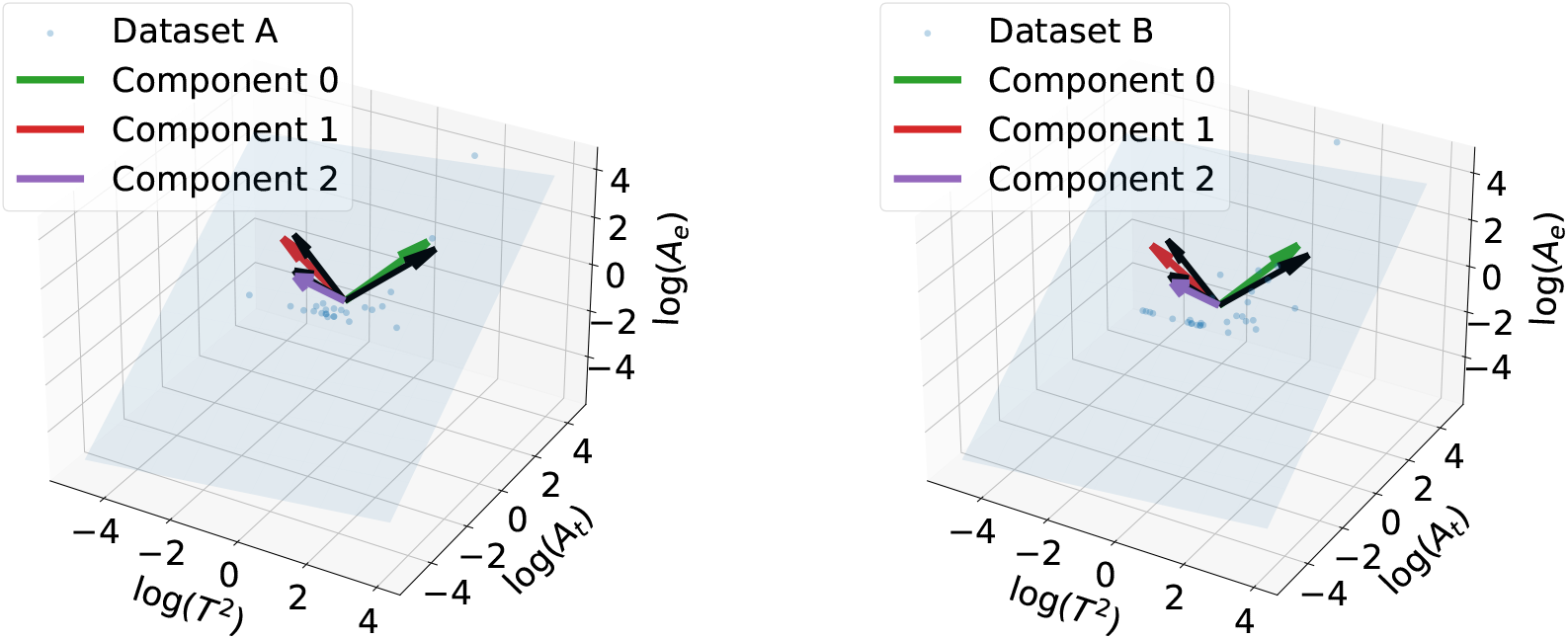
Cortical plane considering different mammalian species datasets and the vectors reconstructed by PCA. The arrows in black represent the principal components predicted by the model, the directions of the natural variables *K, S*, and *I*. To avoid systematic uncertainties from different datasets, we applied the analysis in datasets A (left panel) and B (right panel) separately. The angular difference between theoretical components and the experimental ones are *θ*_diff_ *< 7*^*◦*^. The results are remarkably in agreement with the predictions and consistent between distinct experiments, suggesting that these variables are indeed a natural mathematical language to describe the cortex, encapsulating orthogonal information about the morphology.

**Table 2:** The normalized vectors found by PCA and their respective deviation from theoretical expectation for comparative neuroanatomy. The components are ordered by the Explained Variance Ratio (EVR).

| Dataset | Direction | Theoretical | PCA | $\theta_{\text{diff}}$ | EVR |
| --- | --- | --- | --- | --- | --- |
| A | $\hat{v}$ | $\hat{I} = (0.577, 0.577, 0.577)$ | $(0.490, 0.604, 0.628)$ | $6.0^\circ$ | 0.80 |
| | $\hat{u}$ | $\hat{S} = (-0.802, 0.534, 0.267)$ | $(-0.863, 0.437, 0.253)$ | $6.7^\circ$ | 0.19 |
| | $\hat{w}$ | $\hat{K} = (-0.154, -0.617, 0.771)$ | $(-0.121, -0.666, 0.736)$ | $4.0^\circ$ | 0.01 |
| B | $\hat{v}$ | $\hat{I} = (0.577, 0.577, 0.577)$ | $(0.499, 0.602, 0.623)$ | $5.4^\circ$ | 0.81 |
| | $\hat{u}$ | $\hat{S} = (-0.802, 0.534, 0.267)$ | $(-0.859, 0.439, 0.264)$ | $6.4^\circ$ | 0.18 |
| | $\hat{w}$ | $\hat{K} = (-0.154, -0.617, 0.771)$ | $(-0.114, -0.666, 0.736)$ | $4.1^\circ$ | 0.01 |

The combination of both datasets is discussed in Appendix 5, showing that this PCA method can detect subtle systematic differences between different datasets and underscores the need to properly harmonize said datasets before performing joint analyzes.

### 2.3 Simulation of the PCA performance

In order to address the power of the PCA to recover the true principal components vectors in the comparative neuroanatomy datasets, we ran the same analysis using *N*_sim_ = 10^5^ simulated datasets with known least variance PC direction. Figure 3 shows the distribution of the angle *θ*_diff_ measuring the difference between the theoretical direction and the ones reconstructed by the method. We found the deviation between the real 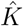 direction simulated and the one reconstructed by PCA was *θ*_diff_ = (6.2^*◦*^ + 2.5^*◦*^ *−* 0.9^*◦*^) comparable to the values found when using real data.

**Figure 3:**
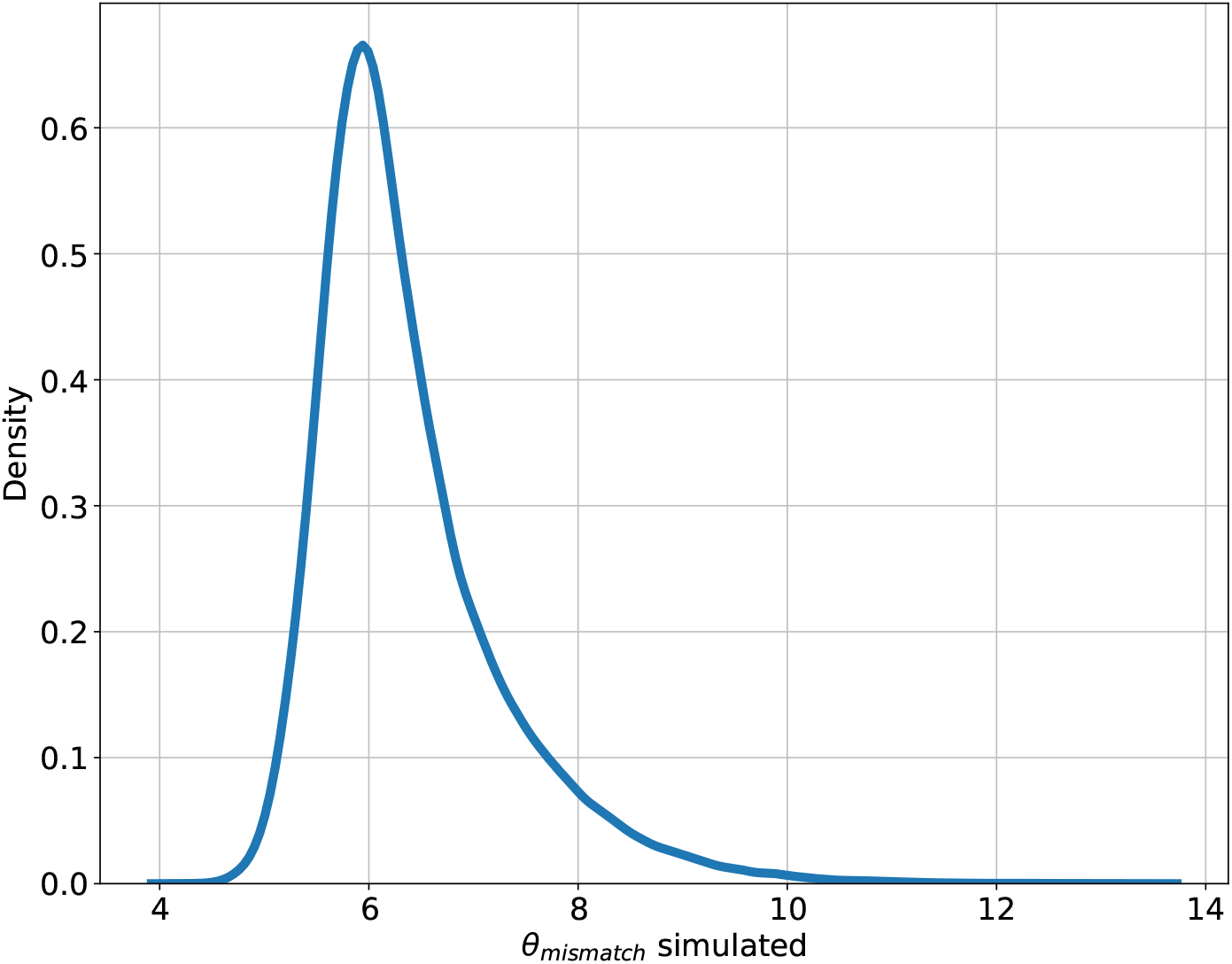
Angular difference *θ*_diff_ between the real principal component 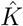 and the one reconstructed by the PCA with the smallest explained variance. The deviation found is *θ*_diff_ = (6.2^*◦*^ + 2.5^*◦*^ *−* 0.9^*◦*^).

### 2.4 Fitting the power-law

An alternative approach to the PCA is the direct determination of the exponents *α, β* and *γ* in Equation 3 using a standard maximum likelihood approach. The analysis starts performing a linear regression of the universal scaling relation in the particular case of lissencephalic mammals results in Figure 4. The angular coefficient 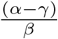 gives priorinformation on the desired parameters.

**Figure 4:**
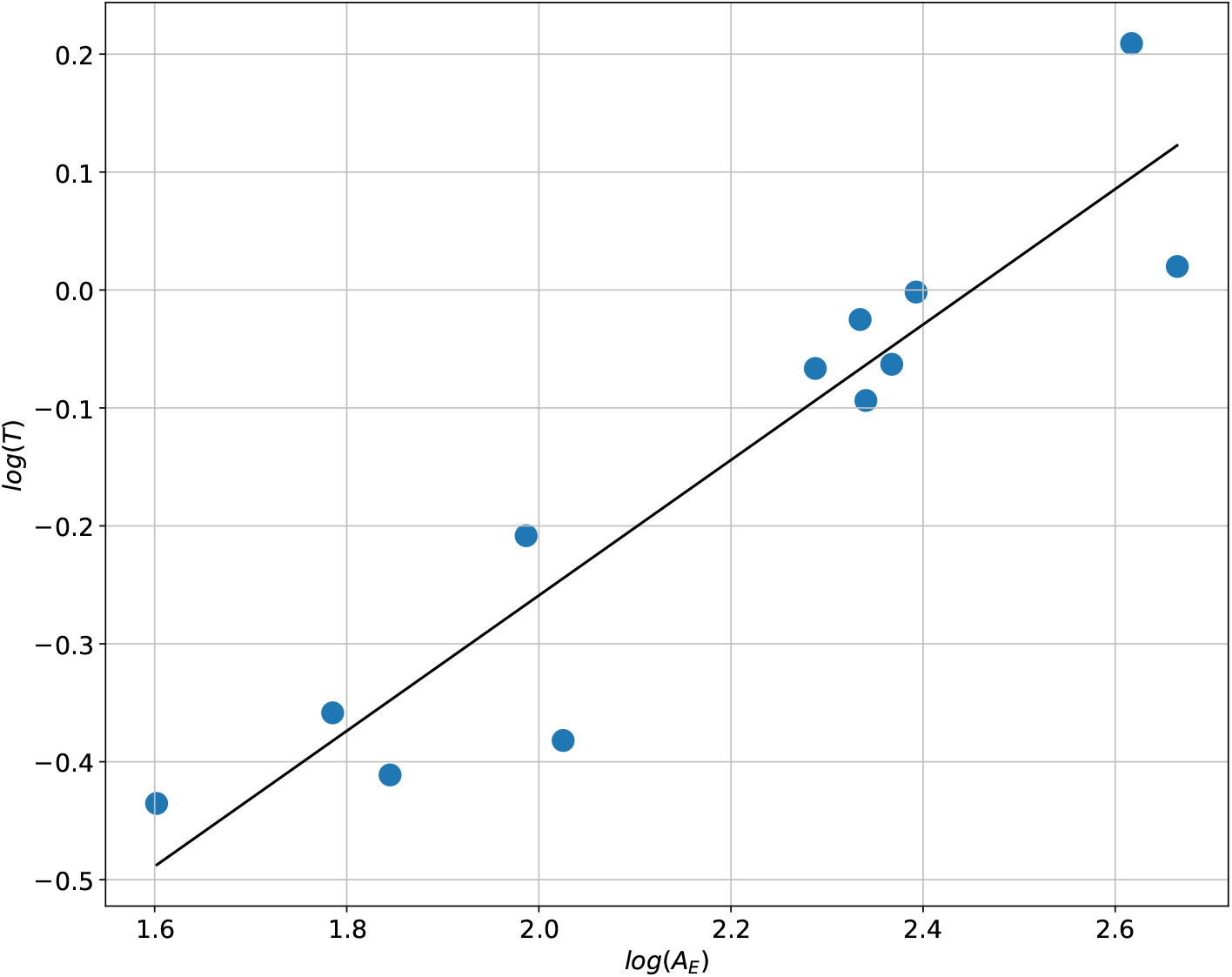
Fitting the universal scaling relation in the particular case of lissencephalic mammals. The angular coefficient 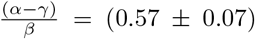 provides a constraint on the possible values for the exponents *α, β* and *γ* in Equation 3 and is in agreement with the theoretical predictions *α =* 5/4, *β =* 1/2 and *γ =* 1.

One may note that the experimental angular coefficient (0.57 *±* 0.07) is compatible with the theoretical predictions ((*α − γ)/β =* (5/4 *−* 1)/(1/2) = 0.5), providing a constraint on the possible values for the exponents. Using this constraint, it is possible to fit the data from gyrencefalic species. Here the human MRI dataset was used because it has significantly more data. Figure 5 shows the log-likelihood as function of the exponents *α* and *β*.

**Figure 5:**
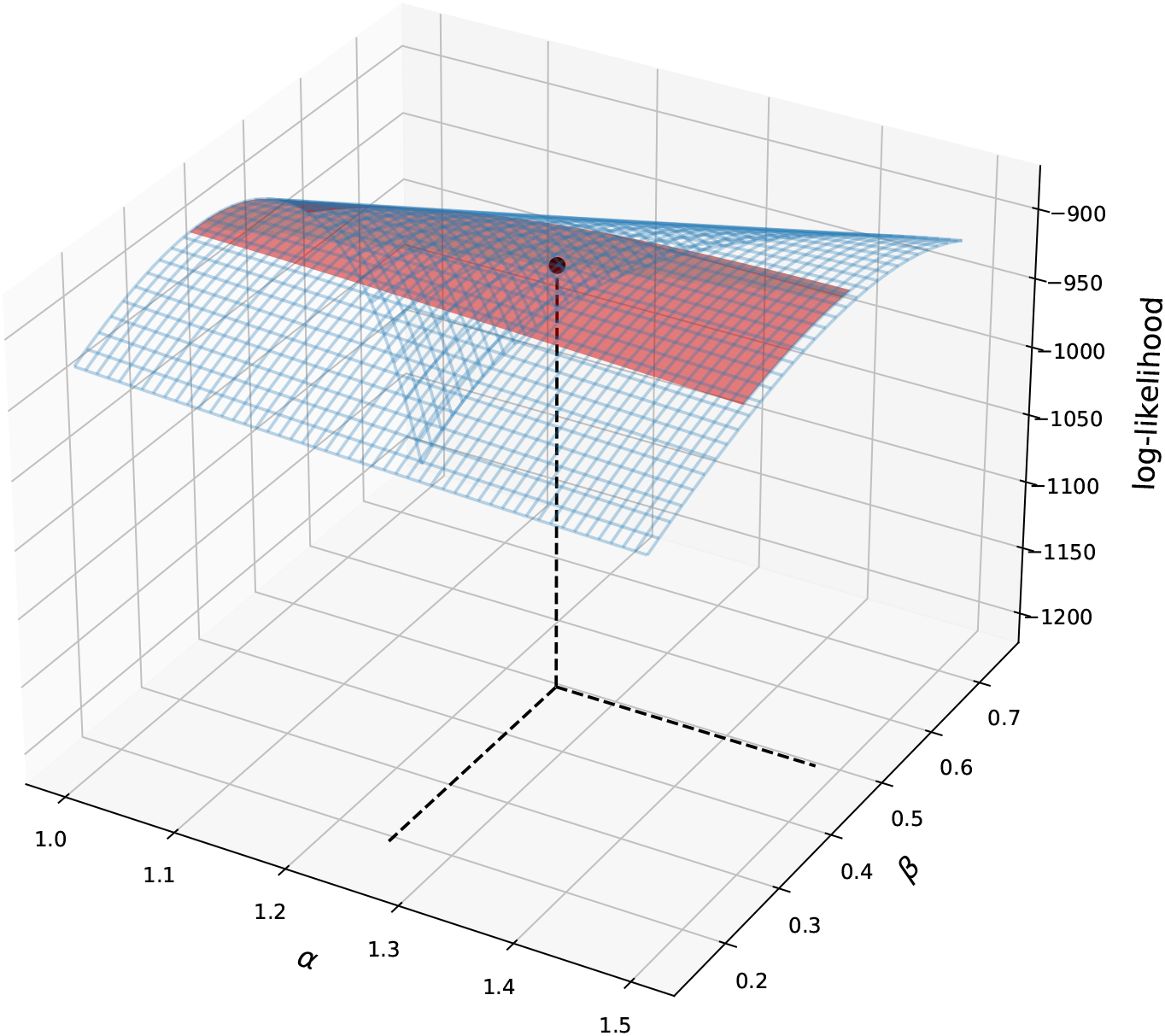
Log-likelihood function with the constraint established by lissencephalic data. Instead of a function peaked around a localized region, it is observed a large range of equivalent peaks. The maximum of the likelihood function with 1 *σ c*onfidence region represented in red. Remarkably, the theoretical expectation (black dot) is in agreement with the values obtained by the regression.

## 3 Discussion

There is already some evidence that the cortical surface behaves as an elastic crumpled surface [22]. Within the framework of this theory, a set of orthogonal morphological variables {*K, S, I}* was proposed as a more natural way of expressing a given cortical shape in the {log *T*^2^, log *A*_*T*_, log *A*_*E*_} morphometric space [38]. The result presented here show that the directions defined by these new variables largely coincide with the principal components of variance for comparative neuroanatomy data of several dozen mammalian species from two independent datasets. Specifically, as expected, the 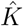 is very close to being the component with the least variance, reflecting the confining of data points at or close to the theoretical cortical plane. The new variables are thus shown to be almost uncorrelated with one another and can be properly regarded as a set of both natural and independent morphometrics, greatly enhancing their potential analytical usefulness.

An additional result derived from the comparative neuroanatomy dataset suggested by the analysis, and further discussed in Appendix 5, is the possible presence of subtle systematic differences between datasets A and B. This underscores the need to properly harmonize this data before performing joint analyzes and may explain the discrepancy between the observed *α*_exp_ = 1.31 *±* 0.01 and the theoretical prediction obtained by [22].

In the human dataset we also observe a remarkable agreement between the directions 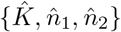, obtained by further imposing the single species constraint of restricted cortical thickness variation, and the principal components of variance. Again, the 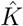 is very close to being the component with the least variance, reflecting the confining of data points at or close to the theoretical cortical plane. This result suggests that the apparent correlation between *S* and *I* in humans is due to the fact that the subjects belong to the same species, a constraint not taken into account when deriving these variables. More than that, in terms of practical application, it means that although the average cortical thickness is nearly invariant within a species and could be used as a biomarker for structural changes in the cortex, it is not an optimal one, as this constraint is evolutionarily contingent. Thus, even for human data, it is valuable to express morphology in terms of the *K, S* and *I v*ariables, as suggested in [17, 41], as these variable independently capture most of the global morphological variation in cortex. This approach contrasts with previous studies such as [4, 6, 8], and may in the end produce more robust biomarkers.

The degree of agreement between morphological variables and principal components was quantified using a simulated dataset, generated with principal components given by the exponents in Equation 1 and the same variance as present in the analyzed dataset. It was found that the deviation between the real 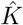 direction simulated and the one reconstructed by PCA was *θ*_diff_ = (6.2^*◦*^ + 2.5^*◦*^ *−* 0.9^*◦*^) (see Figure 3), a value consistent with the one observed in the experimental data.

A deeper explanation of the systematic discrepancy observed between the principal components, the theoretically predicted vectors, and those obtained using the PCA is given by the result of the maximum likelihood approach. The PCA cannot reconstruct the vectorial components with the required precision, reflecting exactly on the distribution shown in Figure 3. The explanation for this impossibility becomes clearer when one realizes that the PCA is a way to reconstruct all exponents in Equation 1 and 3. As shown in Figure 5, instead of a function peaked around a localized region, one obtains a large range of equivalent peaks that, although it is compatible with the theoretical values, fitting these values with high precision is not possible. However, the fact that the all exponents of the power-law were in agreement with the observation is an astonishing new positive evidence in favor of the crumpled cortex model.

The model originally proposed by Mota & Herculano-Houzel suggests that the cortical morphology approximates a real fractal with fractal dimension *f*_dim_ = 2*α*. Different aspects of this model were tested and found to be in agreement with a series of experimental results. Taken together, these results help determine the regime of validity for this model, by mapping the cases in which the overall morphology of the cortical surface can be modeled using this theoretical framework, opening the possibility of biologically interpreting morphological phenomena as cortical folding patterns or even creating reliable simulations of the cortical structure and geometry. The biological interpretation of the cortical folding provided by the Mota & Herculano-Houzel model is a profound statement about its origins. Instead of being a new adaptive property specific to some clades, cortical gyrification is a consequence of the thermodynamic equilibrium of the cortical surface, a mechanism that must be universal to all mammals. This universality is reflected in the scaling of self-similar cortical morphology and its restriction to the cortical plane. Essentially, the difference between a gyrencefalic and a lissencefalic developmental end-states is simply the rate of increase of *A*_*T*_ relative to *T*^2^. The cellular signaling mechanism and genetics regulate the cortical development processes that will result or not in a crumpled surface according to the evolution of each clade and how is the relation between *A*_*T*_ and *T* growing rates. But ultimately, the final surface will reach thermodynamic equilibrium and will result in a self-similar structure with fractal dimension *f*_dim_ = 2*α*

*T*his work makes an addition to the empirical evidence corroborating the crumpled cortex model, showing that not only the principal components of cortical morphological variables are in agreement with the theoretical prediction but also it suggests that the independent set of morphological variables *K, S*, and *I* are indeed a natural mathematical language to describe the cortex, encapsulating orthogonal information about the morphology. Also, by finding the direction 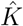 very close to being the component with the least variance in multiple datasets from different mammals, we observe the cortical plane as evidence of a conserved physical mechanism leading the cortical folding. Furthermore, this study shows the possibility of understanding cortical morphology using a unique mathematical description, advocating in favor of incorporating compared neuroanatomy insights in studies of human diseases. The variables discussed here to describe cortical morphology have an immediate practical application when considering disease discrimination and healthy aging studies. Regarding the global morphometry of the cortex, these variables gives the correct treatment to inherent correlation given by the cortical plane. Finally, we also provided a remarkable new evidence in favor of the crumpled cortex model by verifying that each theoretical exponent of the power-law is supported empirically.

## 4 Methods and Materials

### 4.1 Dataset

The morphological data used in this analysis were all previously published. Human participants and their respective MRI-derived metrics were either published by [40, 41] or from open-access databases, as AHEAD [1] and AOMIC (PIOP01 and PIOP02) [30] (human subjects are described in Table 3). The total number of human subjects MRI included in this work is 2456 healthy controls (CTL) from 4 to 96 years old.

**Table 3:** Summary of the subjects used in the human MRI dataset.

| Datasets | Diagnostic | N (F) | Age (Age range) [years] |
| --- | --- | --- | --- |
| AHEAD [1] | CTL | 100 (56) | 42 ± 19 (24; 76) |
| AOMIC PIOP01 [30] | CTL | 208 (120) | 22 ± 1.8 (18; 26) |
| AOMIC PIOP02 [30] | CTL | 224 (128) | 22 ± 1.8 (18; 26) |
| HCP900r [9] | CTL | 881 (494) | 29 ± 3.6 (22; +36) |
| IXI-Guy's | CTL | 314 (175) | 51 ± 16 (20; 86) |
| IXI-HH | CTL | 181 (94) | 47 ± 17 (20; 82) |
| IXI-IOP | CTL | 68 (44) | 42 ± 17 (20; 86) |
| NKI [25] | CTL | 168 (68) | 34 ± 19 (4; 85) |
| OASIS [19] | CTL | 312 (196) | 45 ± 24 (18; 94) |
| **TOTAL** | **CTL** | **2456 (1762)** | **46 ± 23 (4; 96)** |

The structural images from AHEAD, AOMIC ID1000, PIOP1, and PIOP2 were processed in FreeSurfer v6.0.0 [7] without manual intervention at the surfaces [21]. The FreeSurfer localGI pipeline generates the external surface and calculates each vertex’s local Gyrification Index (localGI) [29]. Values of Cortical Thickness, Total Area, and Exposed Area were extracted with Cortical Folding Analysis Tool [39] for both hemispheres.

In order to perform a joint analysis, the morphological variables were harmonized following the prescription given in [3]. The data harmonization procedure is based on the time evolution of the basic morphological variables *T, A*_*T*_, or *A*_*E*_, modeled as an exponential decay with the age *t*_age_. A joint linear fit was performed in the log-linear scale, maintaining the same angular coefficient among all different samples but different linear coefficients. The linear coefficients were then subtracted to harmonize all datasets.

The time evolution of the morphological variables itself can significantly impact the PCA results and the posterior interpretation. To avoid this problem, we used two techniques: (i) restrict the analysis to the age range between 20-40 years (adulthood) as it would be the best representation of a stationary fully developed cortex; (ii) take advantage of the harmonization linear regression, removing the age dependence using each subject age and the angular coefficient fitted, a procedure called here deageing. The latter allows to perform the analysis to the full age range (20-60 years) in which the model was experimentally validated [3]. In Appendix 5, one finds the replication of the analysis for all age ranges on the dataset (with and without the deageing).

Regarding the mammalian dataset from multiple clades, this work used the extracted morphological variables made available by the authors at [22]. This dataset also presents heterogeneity in acquisition, compiling Mota & Herculano-Houzel’s group data (referred here as dataset A) [10, 11, 16, 24, 27, 35] with previously published cortical morphological measurements (dataset B) [2, 5, 12, 20, 26]. The details about each acquisition protocol can be found in the original publications. This variability can also significantly impact the PCA results due to systematic uncertainties from different experiments. To avoid this problem, we applied the analysis using datasets A and B separately. In Appendix 5, one finds the replication of the analysis for the datasets combined.

#### Principal Component Analysis

Principal Component Analysis (PCA) is a standard procedure in multivariate data analysis, usually used to dimensionality reduction in a dataset [14]. With a direct analogy to the calculation of a rigid body principal axes of inertia, it creates a new set of uncorrelated variables, the principal components, solving an eigenvalue/eigenvector problem for the covariance matrix of a distribution or set of points. In this work, the PCA was performed numerically using Python SciPy library [36]. Across this article, the vectors 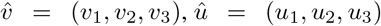 and 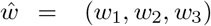 represent the normalized principal components ordered by the size of explained variance.

Considering the {log *T*^2^, log *A*_*T*_, log *A*_*E*_} space, the universal scaling law strongly suggests that the vector 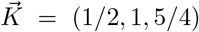 (Equation 2), must be the principal component with the lowest explained variance. It is a near-invariant quantity obtained by isolating *k i*n Equation 1, thus defining the theoretical cortical plane, and in its initial formulation it was associated with conserved viscoelastic properties of cerebral cortical matter (*k ∝* (*f*_*S*_ +*p*_CSF_)^1*/*2^ [15,22]). For humans, for whom in addition to a nearly invariant *K o*ne also finds that the average cortical thickness *T v*aries little [41], one would expect only small variations along the log *T*^2^ direction. Thus, the component with the most variance should lie along the 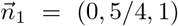 direction. The remaining PC is directly found by the cross product 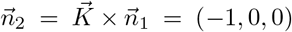. The normalized vectors that will be used in comparison with the PCA are denote here as 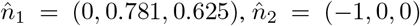, and 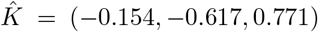.

For the comparative neuroanatomy datasets, there is substantial variation in cortical thickness and areas. Following the interpretation in [38], we predict that the principal components in this case are the normalized directions defined by the independent morphological variables *K*, S and *I* given by *Î* = (0.577, 0.577, 0.577), *Ŝ* = (−0.802, 0.534, 0.267), and 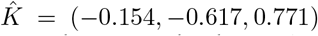. If the variables {*K, S, I}* are actually a natural set of morphological variables, capturing different and orthogonal aspects of the cortex, their respective vectors should be the principal components in the {log *T*^2^, log *A*_*T*_, log *A*_*E*_} space.

### 4.2 Statistical Significance of the PCA results

To better interpret the results above, we used Monte Carlo simulations as a tool to address the power of the analysis to recover the true principal components’ vectors and address the statistical significance of the results. The data was generated over *N*_sim_ = 10^5^ iterations using a Monte Carlo method to sample points near the theoretical cortical plane defined by Equation 2. The number of points generated and their variance was chosen to match the available data on the human MRI dataset. In this simulated dataset, we applied the same PCA procedure used to analyze the real data, calculating the distribution for the angle *θ*_diff_ between the real principal component 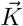 and the component in the PCA with the least explained variance for each simulated dataset.

### 4.3 Fitting the power-law

Assuming a relation in the form of Equation 3, the comparison between PCA least variance component and the theoretical vector is equivalent to perform a joint fit of the exponents *α, β* and *γ* and test if the values are compatible with the theoretical ones, *α =* 5/4, *β =* 1/2 and *γ =* 1. This alternative approach is done using a standard maximum likelihood approach. Firstly, we start fitting the universal scaling relation in the particular case of lissencephalic mammals for which the relation scales as 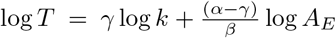. The angular coefficient 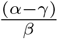 work as a constraint or prior information that can be used fit the data from gyrencefalic species obtaining all parameters.

### 4.4 Data and code availability

All codes and processed datasets are available at https://github.com/bragamello/metaBIO/tree/main/Universal_cortical_rule. AHEAD and AOMIC (PIOP01 and PIOP02) data is available at https://zenodo.org/record/5750619.

## 5 Acknowledgments

This work was supported by Instituto Serrapilheira (grant Serra-1709-16981) and CNPq (PQ 2017 312837/2017-8). We are thankful to the research team and volunteers for participating in this research project. We especially thank Dr. Yujiang Wang for the careful reading and commenting on this manuscript, and for many invaluable suggestions for improvement. The authors have no competing interests.

## Appendix A

**Figure 6:**
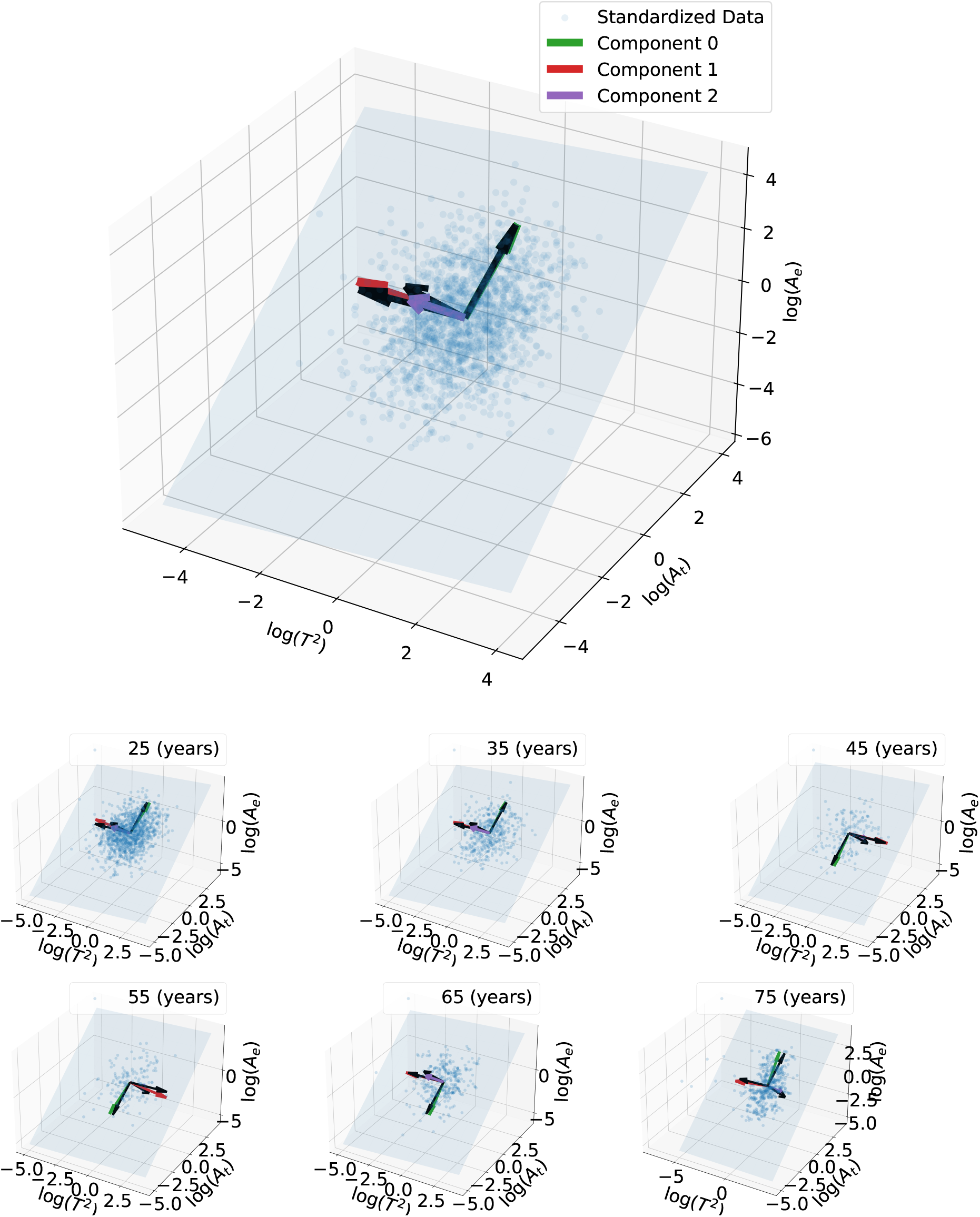
Cortical plane for the human MRI harmonized dataset and the vectors reconstructed by PCA for different age ranges. The arrows in black represent the principal components predicted by the model. An angular difference of 180^*◦*^ on the vectors reflects the ambiguity of selecting the vector normal to the plane.

**Figure 7:**
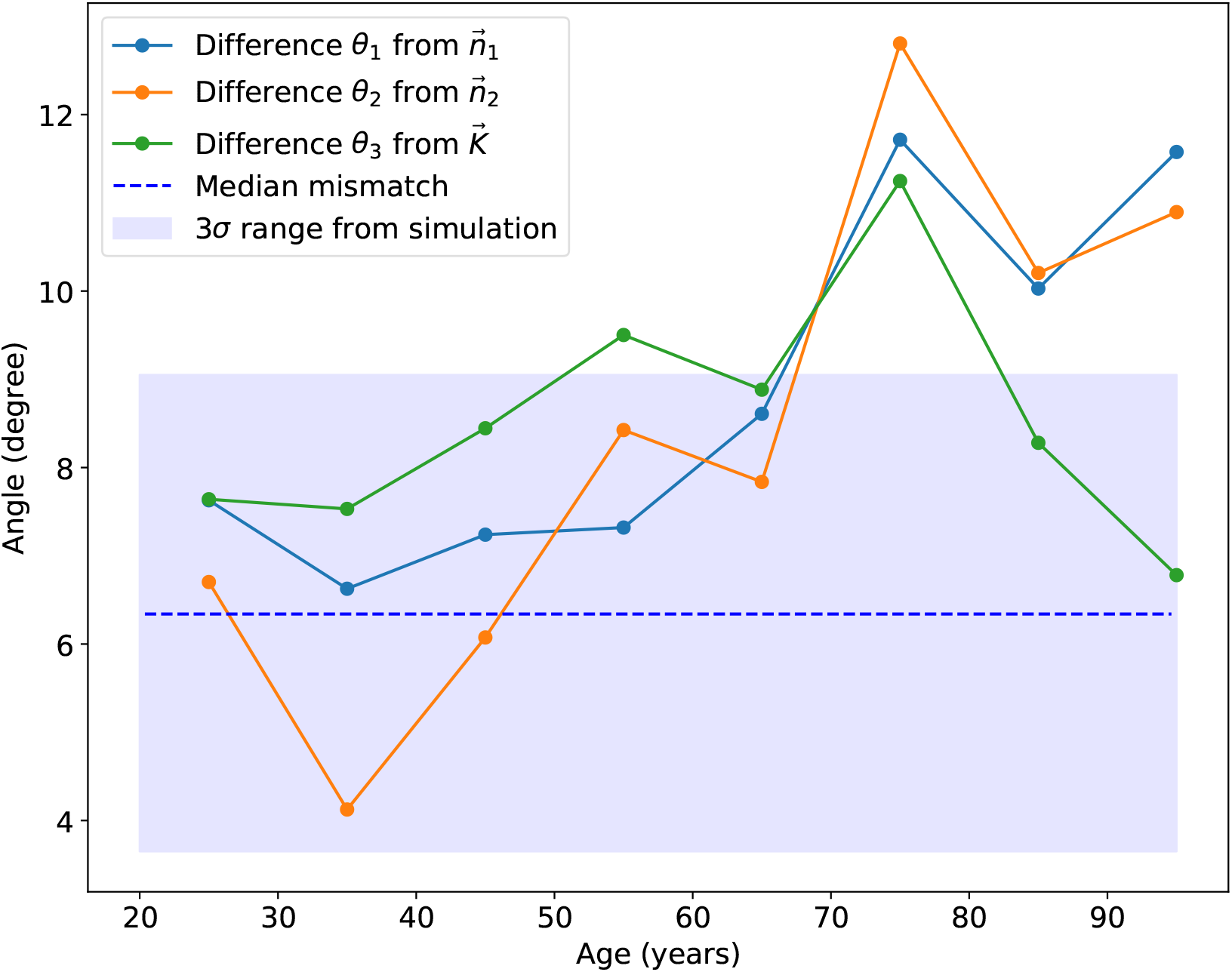
Angular difference between the vectors reconstructed by PCA and the theoretical predictions across healthy aging. The 3*σ r*egion was obtained using the simulations. The results are remarkably in agreement with the predictions when considering ages below 65 years. Above this age, there is a clear change in the directions with the greatest variability. This deviation from the theoretical predictions correlated with age can be further exploited in future works in order to build a theoretical model for the morphology of healthy cortical aging.

**Figure 8:**
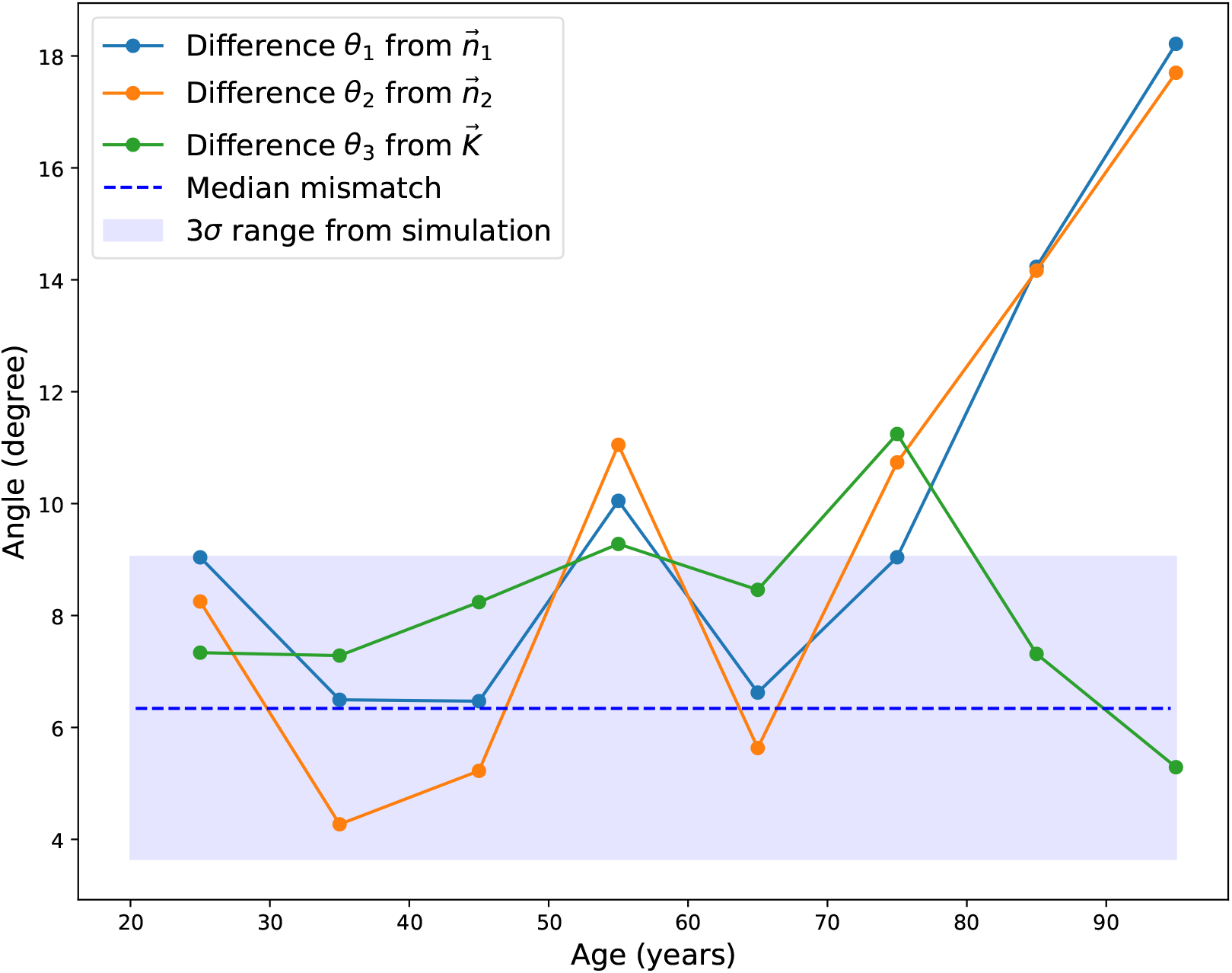
Angular difference between the vectors reconstructed by PCA and the theoretical predictions across healthy aging using the de-aged dataset. The 3*σ r*egion was obtained using the simulations. The results are remarkably in agreement with the predictions when considering ages below 65 years. As expected, without the age dependency, the results are constant until the deviation from the model after 65 years. Above this age, there is a clear change in the directions with the greatest variability. This deviation from the theoretical predictions correlated with age will be further exploited in future works in order to build a theoretical model for the morphology of healthy cortical aging.

## Appendix B

The PCA applied to the combined mammalian dataset, as shown in Figure 9, presents a larger deviation from the theoretical prediction. The vectors found are summarized in table 4.

**Figure 9:**
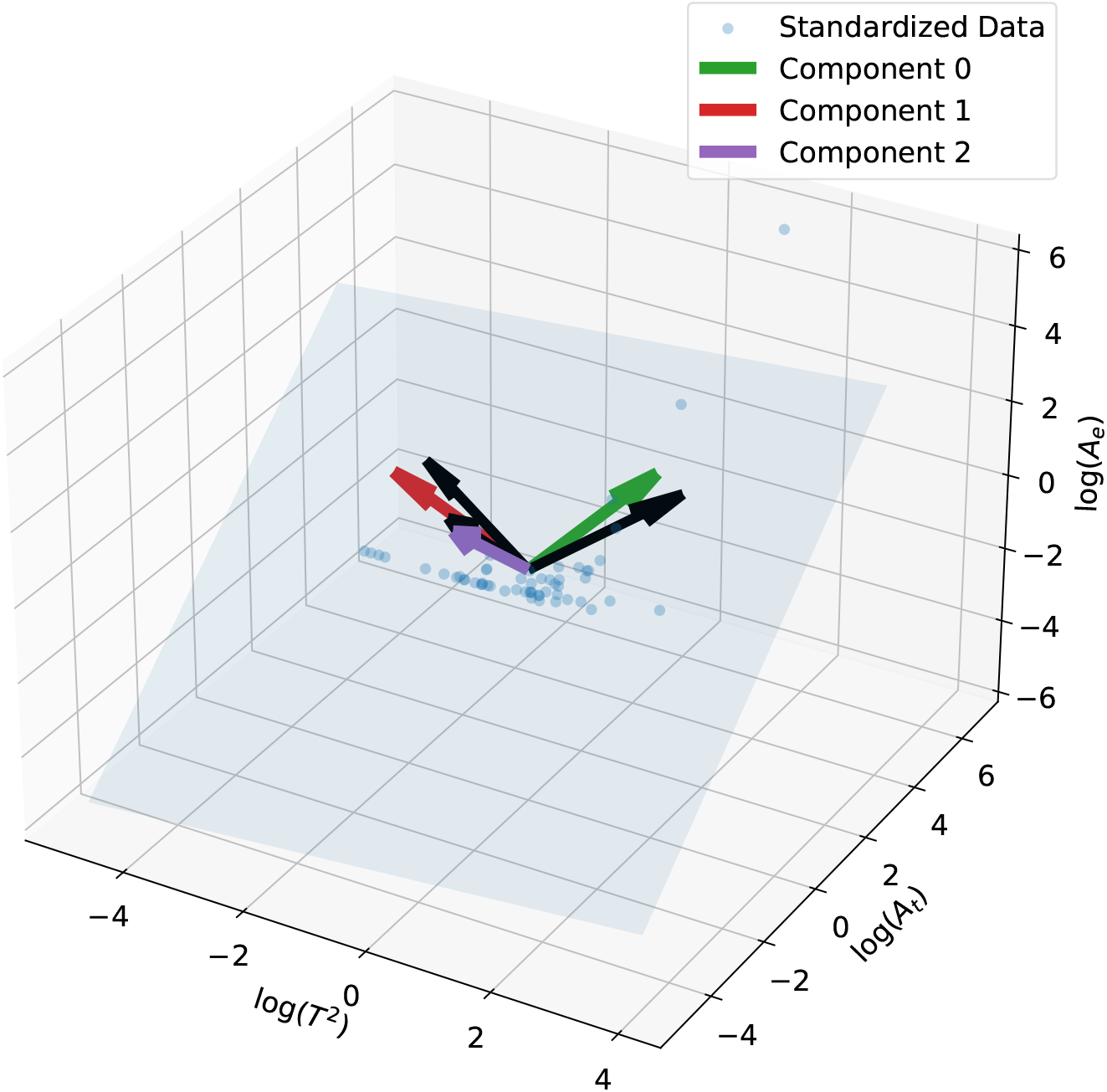
Cortical plane considering different mammalian species datasets and the vectors reconstructed by PCA. The arrows in black represent the principal components predicted by the model, the directions of the natural variables *K, S*, and *I*. In this analysis, datasets A and B were used together. There is an increase in the angular difference between theoretical principal components *Ŝ* and *Î* and the experimental ones. This result is interpreted as indicative of a relative uncounted systematic error between the datasets.

**Table 4:** The normalized vectors found by PCA and their respective deviation from theoretical expectation combining comparative neuroanatomy datasets A and B. The components are ordered by the Explained Variance Ratio (EVR).

| Direction | Theoretical | PCA | $\theta_{\text{diff}}$ | EVR |
| --- | --- | --- | --- | --- |
| $\hat{v}$ | $\hat{I} = (0.577, 0.577, 0.577)$ | $(0.417, 0.632, 0.653)$ | $10.6.0^\circ$ | 0.74 |
| $\hat{u}$ | $\hat{S} = (-0.802, 0.534, 0.267)$ | $(-0.904, 0.364, 0.224)$ | $11.7^\circ$ | 0.25 |
| $\hat{w}$ | $\hat{K} = (-0.154, -0.617, 0.771)$ | $(-0.096, -0.684, 0.723)$ | $5.8^\circ$ | 0.01 |

There is a statistically significant increase in the angular difference from *Ŝ* and *Î*. In comparison with the result presented in the main text, in which both datasets had consistent findings when analyzed separately, there is an indication of a relative unaccounted systematic error between the datasets. This illustrates how this PCA method can be used to detect subtle systematic differences between different datasets and underscores the need to properly harmonize said datasets before performing joint analyses. The suggestion of an unaccounted systematic error between datasets A and B should be further investigated by analyzing the experimental errors of data acquisition, having the potential to elucidate the experimental discrepancy between the reported index *α =* 1.32 and the theoretical 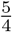.

## Notes

https://github.com/bragamello/metaBIO

https://zenodo.org/record/5750619

### Competing Interest Statement

The authors have declared no competing interest.

### Summary of Updates

This version has an extended introduction to better clarify the theory behind the results. Also, all figures were improved for better visualization

